# Identification of jellyfish neuropeptides that act directly as oocyte maturation inducing hormones

**DOI:** 10.1101/140160

**Authors:** Noriyo Takeda, Yota Kon, Gonzalo Quiroga Artigas, Pascal Lapébie, Carine Barreau, Osamu Koizumi, Takeo Kishimoto, Kazunori Tachibana, Evelyn Houliston, Ryusaku Deguchi

## Abstract

Oocyte meiotic maturation is a critical process for sexually reproducing animals, and its core cytoplasmic regulators are highly conserved between species. In contrast, the few known Maturation Inducing Hormones (MIHs) that act on oocytes to initiate this process have highly variable molecular natures. Using the hydrozoan jellyfish species *Clytia* and *Cladonema*, which undergo oocyte maturation in response to dark-light and light-dark transitions respectively, we deduced from gonad transcriptome data amidated tetrapeptide sequences and found that synthetic peptides could induce maturation of isolated oocytes at nanomolar concentrations. Antibody preabsorption experiments conclusively demonstrated that these W/RPRPamide-related neuropeptides account for endogenous MIH activity produced by isolated gonads. We further showed that the MIH peptides are synthesised by neural-type cells in the gonad, are released following dark-light / light-dark transitions, and probably act on the oocyte surface. They are produced by male as well as female jellyfish and can trigger both sperm and egg release, suggesting a role in spawning coordination. We propose an evolutionary link between hydrozoan MIH and the neuropeptide hormones that regulate reproduction upstream of MIH in bilaterian species.

## Introduction

Fully-grown oocytes maintained within the female gonad are held at first prophase of meiosis until environmental and/or physiological signals initiate cell cycle resumption and oocyte maturation, culminating in release of fertilisation-competent eggs. This process of oocyte maturation is a key feature of animal biology, tightly regulated to optimise reproductive success. It involves biochemical cascades activated within the oocyte that are highly conserved across animal phyla, notably involving the kinases Cdk1 (to achieve entry into first meiotic M phase) and MAP kinase (to orchestrate polar body formation and cytostatic arrest) (Amiel et al., 2009; Von Stetina and Orr-Weaver, 2011; Tachibana et al., 2000; Yamashita et al., 2000). These kinase regulations have been well characterised using biochemically tractable model species, notably frogs and starfish, and knowledge extended using genetic methods to other species including nematodes, drosophila and mammals. Nevertheless information is largely lacking on certain critical steps, and in particular the initial triggering of these cascades in response to the Maturation Inducing Hormones (MIHs), which act locally in the gonad on their receptors in the ovarian oocytes; the only known examples identified at the molecular level are 1-methyladenine released in starfish (Kanatani et al., 1969), steroid hormones in amphibians and fish (Haccard et al., 2012; Nagahama and Yamashita, 2008), and a sperm protein in *Caenorhabditis* (Von Stetina and Orr-Weaver, 2011).

Hydrozoan jellyfish provide excellent models for dissecting the molecular and cellular mechanisms regulating oocyte maturation, which in these animals is triggered by light-dark and/or dark-light transitions. Remarkably, oocyte growth, maturation and release continue to function autonomously in gonads isolated from female jellyfish, implying that all the regulatory components connecting light sensing to spawning are contained within the gonad itself (Amiel et al., 2010; Freeman, 1987; Ikegami et al., 1987). Furthermore, as members of the Cnidaria, a sister clade to the Bilateria, hydrozoan jellyfish can give insight into spawning regulation in early animal ancestors.

In this study we addressed the molecular nature and the cellular origin of MIH using two hydrozoan jellyfish model species *Clytia hemisphaerica* (Fig. 1A) and *Cladonema pacificum* (Fig. 1B). These species are induced to spawn by dark-light or light-dark transitions respectively (Amiel et al., 2010; Deguchi et al., 2005; Houliston et al., 2010). Starting from the hypothesis that hydrozoan MIH might comprise neuropeptides, consistent with size filtration and protease sensitivity experiments (Ikegami et al., 1987), we screened synthetic candidate peptides predicted from gonad transcriptome data by treatment of isolated gonads (spawning assay) or isolated oocytes (MIH assay). We then raised inhibitory antibodies to confirm the presence and activity of the putative peptides in native MIH secreted from gonads in response to light/dark cues, and to characterise the MIH-producing cells and their response to light. We extended our findings by determining the activity of the identified hydrozoan MIH tetrapeptides on males as well as females, and on a selection of other diverse hydrozoan species. A parallel study of *Clytia* gonad light detection revealed that the light-mediated MIH release reported here is dependent on an Opsin photopigment co-expressed in the same population of cells that secretes MIH (Quiroga Artigas et al., 2017). These specialised cells, which have neural-type morphology and characteristics, thus provide a simple and possibly ancestral mechanism to promote synchronous gamete maturation, release and fertilisation.

**Figure 1.**
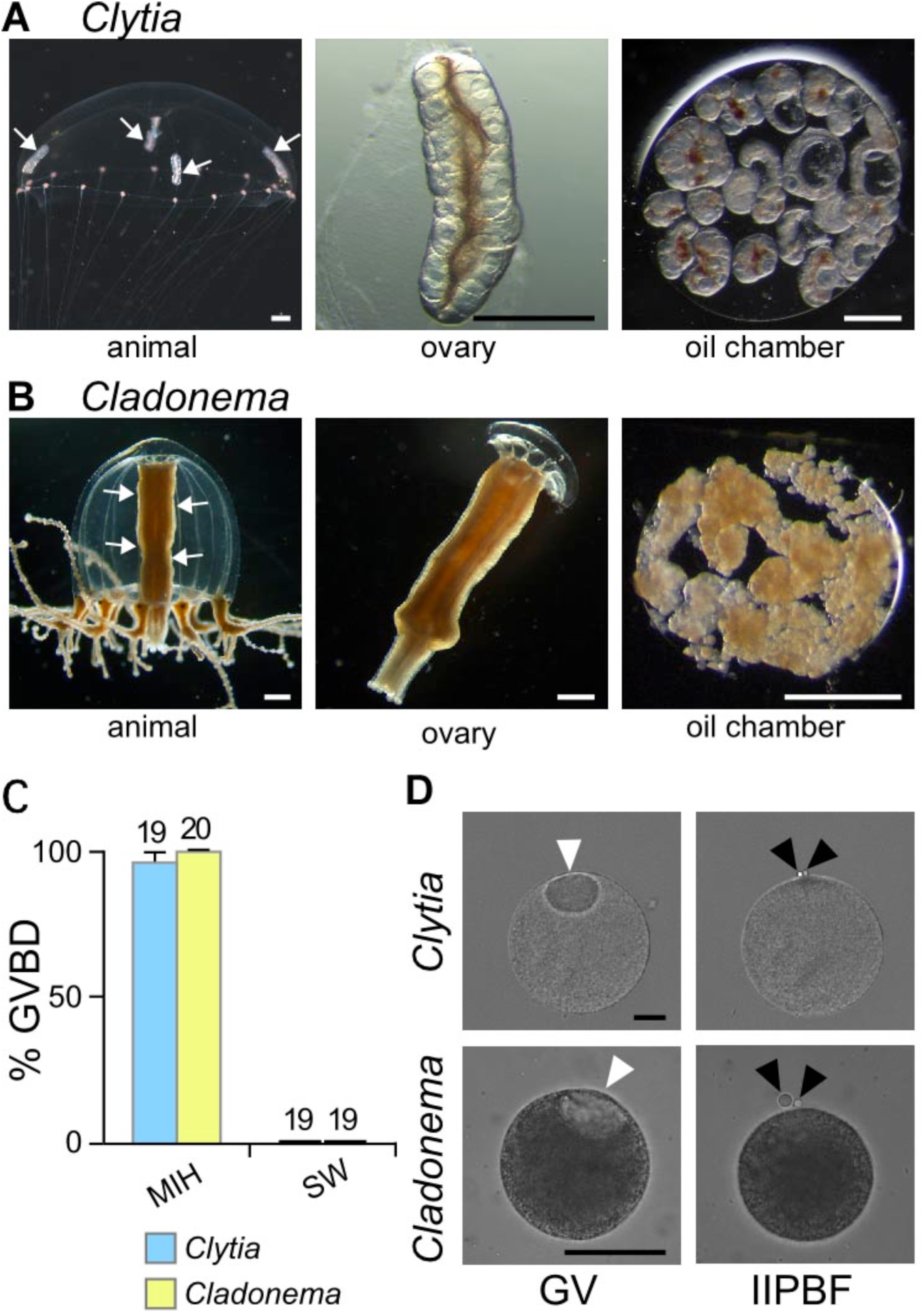
Active MIH is produced by isolated jellyfish gonads. A) *Clytia hemisphaerica* whole female jellyfish (1cm diameter), isolated ovary and a collection of ovaries under oil used to collect MIH. B) Equivalent samples for *Cladonema pacificum.* Arrows point to gonads in A and B. C) GVBD assay on isolated oocytes (number of oocytes marked above each bar) incubated in presence or absence of MIH from the same species. D) Isolated oocytes from each species before (‘GV’ stage) and 2 or 1 h respectively after addition of MIH at the time of 2nd polar body formation (“IIPBF” stage). White arrowheads point to GVs and black arrowheads to polar bodies. Scale bars: 500 *μ*m in A, B and 50 μm in D.

## Results

### *Active MIH can be recovered from isolated* Clytia *and* Cladonema *gonads*

First we demonstrated that true MIH activity can be recovered from small drops of seawater containing isolated ovaries of either *Clytia* or *Cladonema* following the appropriate light transition, as demonstrated previously using other hydrozoan species (Freeman, 1987; Ikegami et al., 1987). Isolated oocytes incubated in endogenous MIH recovered using this method complete efficiently the meiotic maturation process, manifest visually by germinal vesicle breakdown (GVBD) and extrusion of two polar bodies (Fig. 1C,D).

Further characterisation of MIH using *Cladonema* showed that isolated gonad ectoderm, but not endoderm, tissue (see Fig. 1B) could produce active MIH. MIH activity from *Cladonema* gonad ectoderm resisted heat treatment at 100°C for 20 minutes (95% GVBD, n=41), several freeze/thaw cycles (100% GVBD, n=14) and to filtration through a 3000 MW cut-off membrane (90 % GVBD, n=18), consistent with the idea that the active molecule is a small molecule, possibly peptidic (Ikegami et al., 1987).

### MIH candidates identified from transcriptome data

Cnidarians including jellyfish, hydra and sea anemones express many low-molecular-weight neuropeptides showing various bioactivities (Anctil, 2000; Fujisawa, 2008; Takahashi et al., 2008; Takeda et al., 2013). These are synthesised by cleavage of precursor polypeptides and can produce multiple copies of one or more peptides, frequently subject to amidation by conversion of a C-terminal glycine (Grimmelikhuijzen et al., 1996). A previous study found that some synthetic *Hydra* amidated peptides can stimulate spawning when applied to gonads of the jellyfish *Cytaeis uchidae*, the most active being members of the GLWamide family (active at 10^−5^ M minimum concentration) (Takeda et al., 2013). Critically, however, these did not induce meiotic maturation when applied to isolated oocytes, i.e. they did not meet the defining criterion of MIHs. These previous results were not conclusive because of the use of species-heterologous peptides, but suggest that although jellyfish GLWamide peptides do not act as MIHs, they may be involved less directly in spawning regulation.

To identify endogenous species-specific neuropeptides as candidates for MIH from our model species, we first retrieved sequences for 10 potential amidated peptide precursors from a mixed-stage *Clytia* transcriptome (Fig. S1), and then searched for ones specifically expressed in the ectoderm, source of MIH, by evaluating the number of corresponding Illumina Hiseq reads obtained from manually separated ectoderm, endoderm and oocyte gonad tissues (Fig. 2A). In the ectoderm, source of MIH, only 3 putative neuropeptide precursor mRNAs were expressed above background levels, as confirmed by quantitative PCR (Fig. S2). One was a GLWamide precursor, Che-pp11, expressed at moderate levels. Much more highly expressed were Che-pp1 and Che-pp4, both predicted to generate multiple related short (3-6 amino acid) amidated peptides with the C-terminal signature (W or R)-PRP, -PRA -PRG or -PRY.

**Figure 2.**
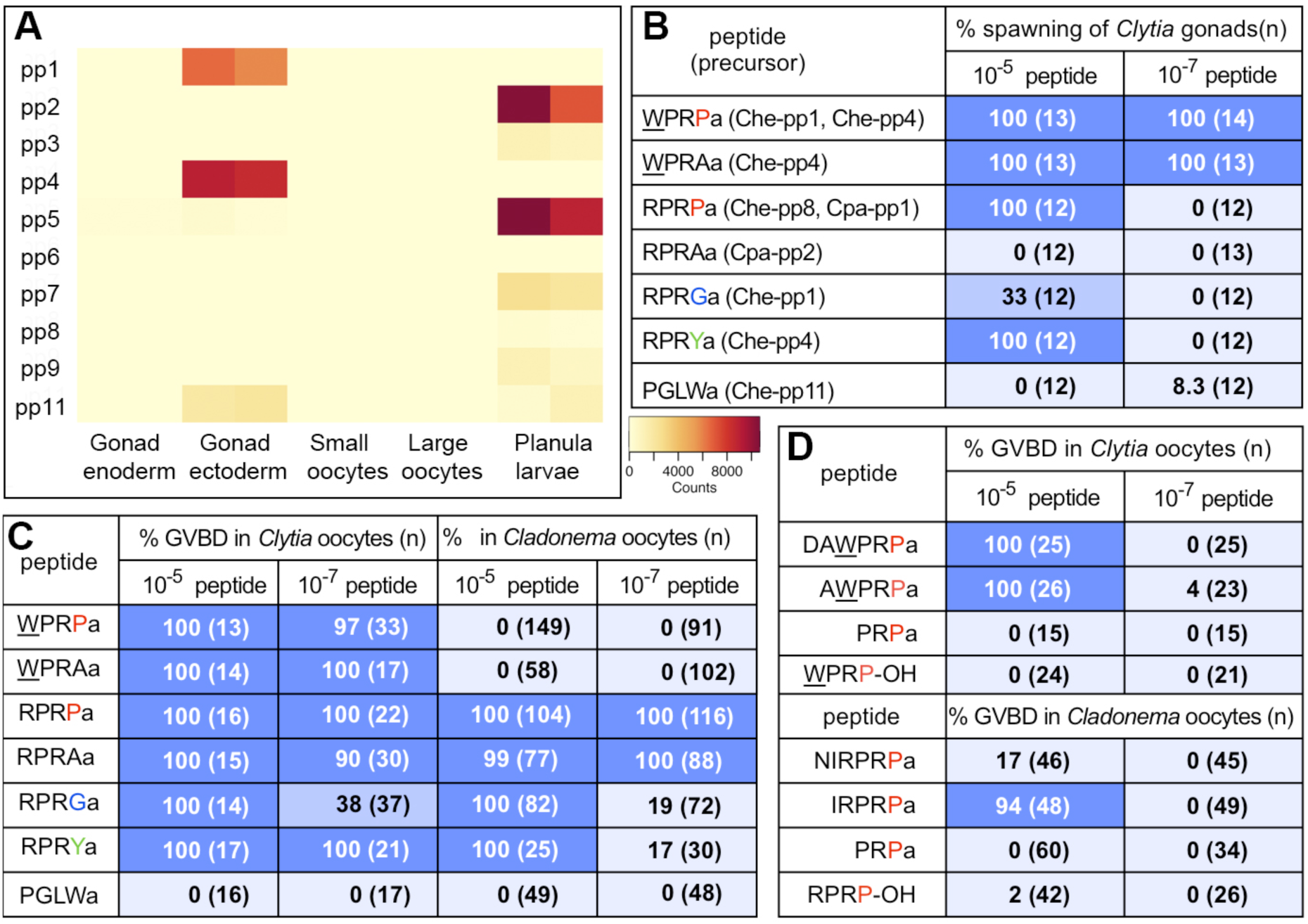
Predicted neuropeptides from the gonad ectoderm have MIH activity. A) Heat map representing the expression of 10 candidate peptide precursor sequences from *Clytia hemisphaerica* in isolated ectoderm, endoderm, small (growing) and large (fully-grown) oocytes from mature female gonads. Illlumina High-seq 50nt reads generated from ectoderm, endoderm and oocyte mRNA were mapped against a *Clytia* reference transcriptome. Data from a sample of 2 day old planula larvae are included for comparison. B) Results of spawning assay on isolated *Clytia* gonads using synthesised amidated tetrapeptides; WPRPamide and WPRAamide, generated from Che-pp1 and Che-pp4 precursors, induced 100% spawning even at 10^−7^ M. C) MIH assay using isolated *Clytia* or *Cladonema* oocytes showing strong MIH activity of related amidated tetrapeptides. D) Synthesised amidated 3, 5 or 6 amino acid peptides, and non-amidated tetrapeptides, show poor MIH activity on isolated *Clytia* and *Cladonema* oocytes.

Potential precursors for both GLWamide (Cpa-pp3) and PRP/Aamides (Cpa-pp1 and Cpa-pp2) were also present amongst 4 sequences identified in a transcriptome assembly from the *Cladonema* manubrium (which includes the gonad; Fig. 1B). Cpa-pp1 contains 1 copy of the RPRP motif while Cpa-pp2 contains multiple copies of RPRA motifs (Fig. S1).

### Potent MIH activity of synthetic W/RPRXamide peptides

As a first screen to select neuropeptides potentially involved in regulating oocyte maturation, we incubated *Clytia* female gonads in synthetic tetrapeptides predicted from Che-pp1, Che-pp4 and Che-pp11 precursors at 10^−5^ M or 10^−7^ M (Fig. 2B). We uncovered preferential and potent activity for the WPRPamide and WPRAamide tetrapeptides, which consistently provoked oocyte maturation and release from the gonad at 10^−7^ M, while the related RPRGamide and RPRYamide were also active but only at 10^−5^ M. RPRPamide, a predicted product of Cpa-pp1 and also of Che-pp8, a precursor not expressed in the *Clytia* gonad, was also active in this screen at 10^−5^ M. In contrast PGLWamide, potentially generated from Che-pp11, did not affect the gonads at either concentration. This result placed WPRP/Aamide-related peptides as the best candidates for jellyfish MIH. We then performed direct MIH activity assay, i.e. treatment of isolated oocytes with the candidate peptides. For *Clytia* oocytes we detected potent MIH activity (as assessed by oocyte Germinal Vesicle breakdown; GVBD; Fig. 2C; Fig. S3) for W/RPRP/Aamide and RPRY/Gamide tetrapeptides, but not for PGLWamide. RPRAamide was more active in triggering GVBD when added to isolated oocytes than to intact gonads, perhaps because of poor permeability through the gonad ectoderm. For *Cladonema* oocytes, RPRP/Aamides showed very potent MIH activity, and the RPRG/Yamides were also active at higher concentrations, but WPRP/Aamides were not active (Fig. 2C, S3). We also tested, on oocytes of both species, pentapeptides and hexapeptides that might theoretically be generated from the Che-pp1 and Cpa-pp1 precursors, but these had much lower MIH activity than the tetrapeptides, while the tripeptide PRPamide and tetrapeptides lacking amidation were inactive (Fig. 2D, S3). The response of *Clytia* or *Cladonema* isolated oocytes elicited by synthetic W/RPRP/A/Yamides mirrored very closely that of endogenous MIH, proceeding through the events of oocyte maturation with normal timing following GVBD, advanced by 15-20mins (*Clytia*) or 10 minutes *(Cladonema)* compared to light /dark induced spawning of gonads (Fig. S4A-D). The resultant mature eggs could be fertilised and develop into normal planula larvae (Fig. S4E,F).

### W/RPRPamides account for endogenous MIH activity

We demonstrated that W/RPRPamide and/or W/RPRAamide peptides are responsible for endogenous MIH activity in *Clytia* and *Cladonema* by use of inhibitory affinity purified antibodies generated to recognise the PRPamide and PRAamide motifs (as determined by ELISA assay; Fig. 3A,B). These antibodies were able to inhibit specifically the MIH activity of the targeted peptides (Fig. 3C,D). Conclusively, pre-incubation of endogenous MIH obtained from *Clytia* or *Cladonema* gonads with anti-PRPamide antibody for 30 minutes completely blocked its ability to induce GVBD in isolated oocytes. Pre-incubation with the anti-PRAamide antibody slightly reduced MIH activity but not significantly compared to a control IgG (Fig. 3E).

**Figure 3.**
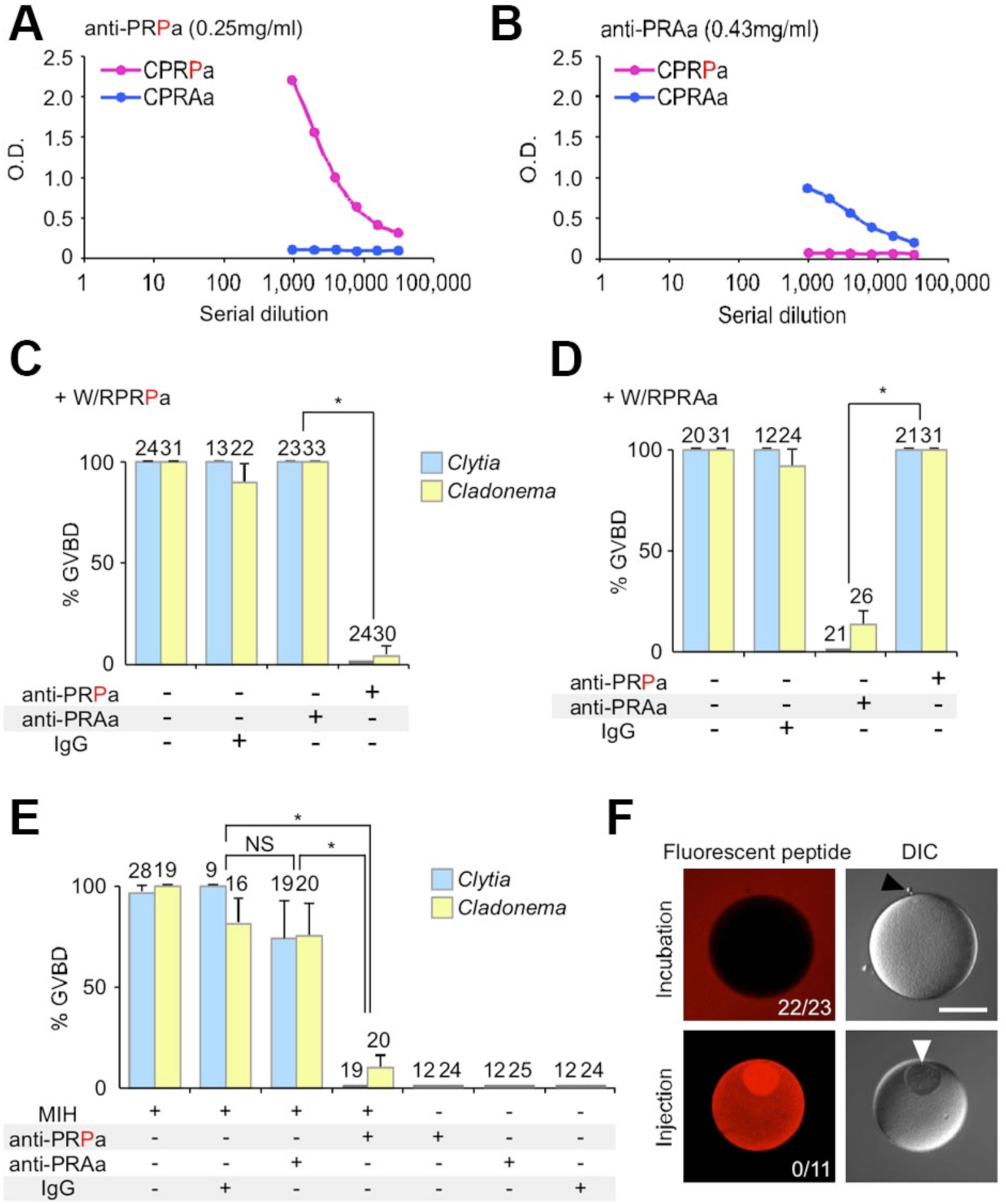
Figure 3. Antibody inhibition shows that PRPamides are the active component of MIH. (A)ELISA assay demonstrating that the anti-PRPamide antibody binds PRPamide but not PRAamide tetrapeptides. B) Reciprocal specificity demonstrated for the anti-PRAamide antibody. C-E) Inhibition experiments in which either anti-PRPamide or anti-PRAamide antibody was pre-incubated with W/RPRPamide, W/RPRAamide or natural MIH prior to the MIH assay (number of oocytes tested above each bar). Oocyte maturation induced by WPRPamide (*Clytia*) or RPRPamide (*Cladonema*) was inhibited by anti-PRPamide but not anti-PRAamide antibodies, while PRAamide activity was specifically neutralised by anti-PRAamide antibodies. The activity of endogenous MIH produced by either *Clytia* or *Cladonema* gonads was inhibited by anti-PRPamide antibody. Inhibition by the anti-PRAamide antibody was not statistically significant (Student’s *t*-tests; asterisk: *P* < 0.01; NS: *P* > 0.05). F) Confocal images of *Clytia* oocytes which underwent GVBD following incubation in TAMRA-WPRPamide (top), but not following injection of TAMRA-WPRPa (bottom). Numbers indicate GVBD/oocytes tested. A black arrowhead points to polar bodies and a white arrowhead to the GV. Scale bars: 100 *μ*m. In C-F, oocytes that did not mature underwent normal GVBD induced by subsequent addition of excess neuropeptides (10^−5^–10^−7^ M WPRPamide for *Clytia*; 10^−7^ M RPRP/Aamide for *Cladonema*).

Taken together these experiments demonstrate that WPRPamide and RPRPamide are the active components of endogenous MIH in *Clytia* and *Cladonema* respectively, responsible for triggering oocyte meiotic maturation. Other related peptides including RPRYamide, RPRGamide, WPRAamide (*Clytia*) and RPRAamide (*Cladonema*) also probably contribute to MIH. These peptides almost certainly act at the oocyte surface rather than intracellularly, since fluorescent (TAMRA-labelled) WPRPamide microinjected into *Clytia* oocytes, unlike externally applied TAMRA-WPRPamide, did not induce GVBD (Fig. 3F).

### MIH is produced by neurosecretory cells in the gonad ectoderm

Single and double-fluorescence *in situ* hybridisation showed that the *Clytia* MIH precursors Che-pp1 and Che-pp4 are co-expressed in a distinctive population of scattered cells in the gonad ectoderm in males and females (Fig. 4A,B; Fig. 5E). Similarly in *Cladonema*, the predicted RPRPamide precursor Cpa-pp1 was expressed in scattered cells in the manubrium ectoderm, which covers the female or male germ cells (Fig. 4A; Fig. 5A,E).

**Figure 4.**
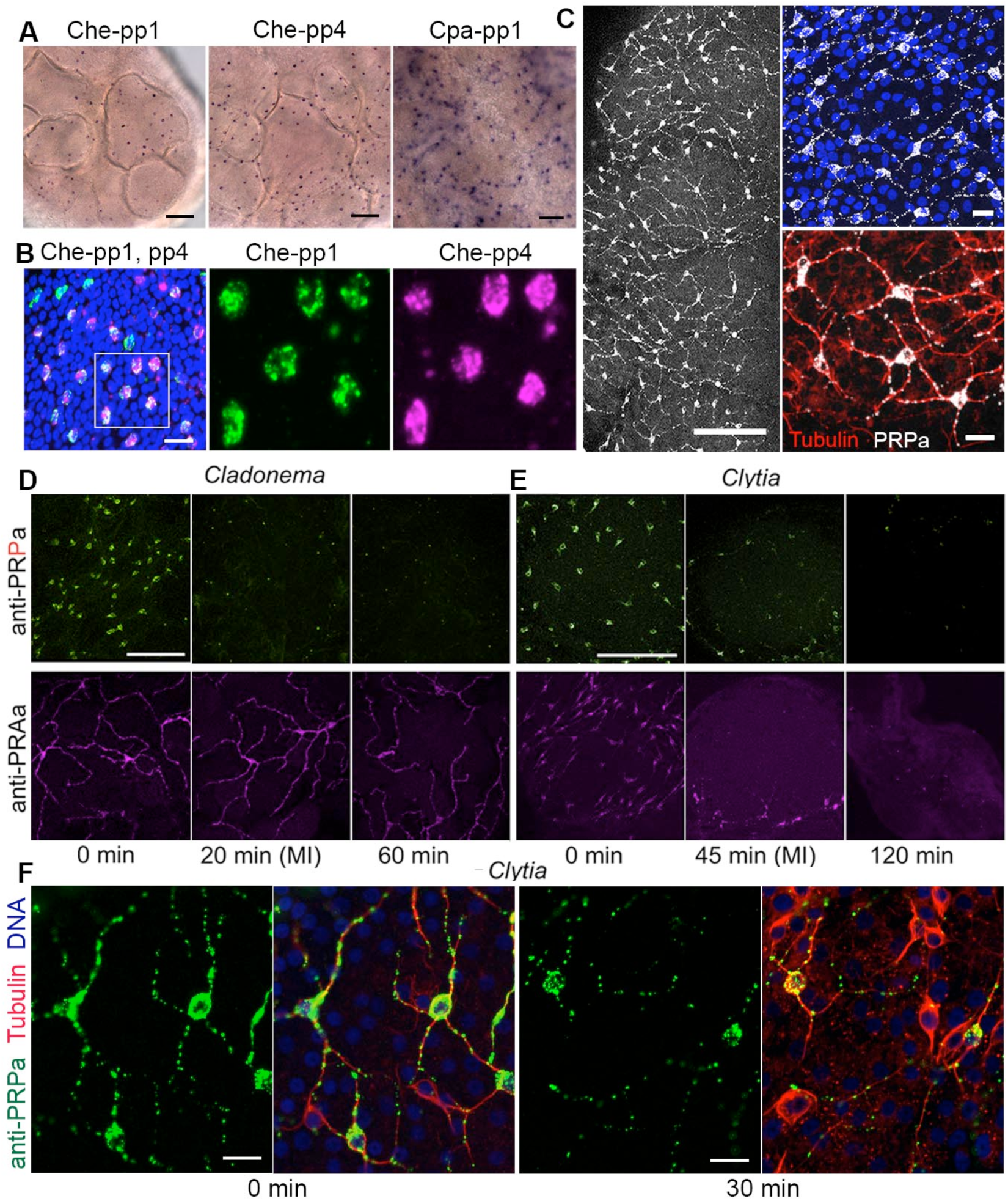
MIH is generated by gonad ectoderm cells with neural characteristics. *In situ* hybridisation detection of Che-pp1 and Che-pp4 mRNAs in *Clytia* (left, center) and of Cpa-pp1 in *Cladonema* (right) in scattered ectoderm cells of female gonads. B) Double fluorescence *in situ* hybridisation reveals co-expression of Che-pp1 (green) and Che-pp4 (magenta); nuclei (Hoechst) in blue. Single channels are shown for the outlined zone in the left image. C) Immunofluorescence of *Clytia* female gonads showing the neural-type morphology of MIH-producing cells, which are characterised by two or more long projections containing microtubule bundles. Staining with anti-PRPamide (white), anti-tubulin (red) and Hoechst (blue). D) Loss of anti-PRPamide staining from *Cladonema* gonads during dark-induced meiotic maturation (MI= first meiotic M phase). The distinct anti-PRAamide stained cells were not obviously affected. E) Equivalent experiment in *Clytia*, in which the two antibodies decorate distinct cell populations- See Figure S5. F) High magnification images of PRPa stained cells in the ectoderm of isolated *Clytia* gonads before (“0 minutes”) or 30 minutes after light exposure. The dots of anti-PRPa staining (green) presumably represent peptide-filled vesicles, which become less abundant in both cell bodies and in the microtubule-rich projections (anti-tubulin staining in red, Hoechst in blue). All fluorescence panels are confocal images. Scale bars: 50 *μ*m in A; 20 *μ*m in B; 100 *μ*m in C (left), D, E; 10 *μ*m in C (right), F.

Immunofluorescence with the anti-PRPamide and anti-PRAamide antibodies in both species revealed that the expressing cells have a morphology typical of cnidarian neural cells, comprising a small cell body and two or more long projections (David 1973), and characterised by the presence of bundles of stable microtubules (Fig. 4C,F). A parallel study further revealed that in *Clytia* these MIH-secreting cells express an opsin photoprotein with an essential function in oocyte maturation and spawning (Quiroga Artigas et al., 2017). Given their neural-type morphology, their photosensory function and their key role in regulating sexual reproduction via neuropeptide hormone production, we propose that these cells have both a sensory and neurosecretory nature. Scattered endocrine cells with both sensory and neurosecretory features are a feature of cnidarians (Hartenstein, 2006). Further, the distribution and organisation of the gonad MIH-producing cells in *Clytia* and *Cladonema* are suggestive of the neural nets that characterise cnidarian nervous systems (Koizumi, 2016; Bosch et al., 2017; Dupre and Yoste, 2017). We could not confirm from our immunofluorescence analyses any direct connections between neighbouring cells. Future electron microscopy or calcium imaging techniques (Gründer and Assmann, 2015; Dupre and Yoste, 2017) to identify synapses or action potential transmission could resolve whether these scattered endocrine cells are integrated within a neural network. In intact *Clytia* jellyfish, both immunofluorescence and *in situ* hybridisation (Fig. 5B-D, S5A) revealed the presence of MIH peptides and their precursors at additional sites associates with neural systems: the manubrium (mouth), tentacles and the nerve ring that runs around the bell rim (Koizumi et al., 2015), as well as along the radial canals. This suggests that PRPamide family neuropeptides have other functions in the jellyfish in addition to regulating spawning. Neuropeptides can have both neural and endocrine functions in cnidarians, and are thought to have functioned in epithelial cells in primitive metazoans prior to nervous system evolution (Hartenstein, 2006; Bosch et al., 2017).

**Figure 5.**
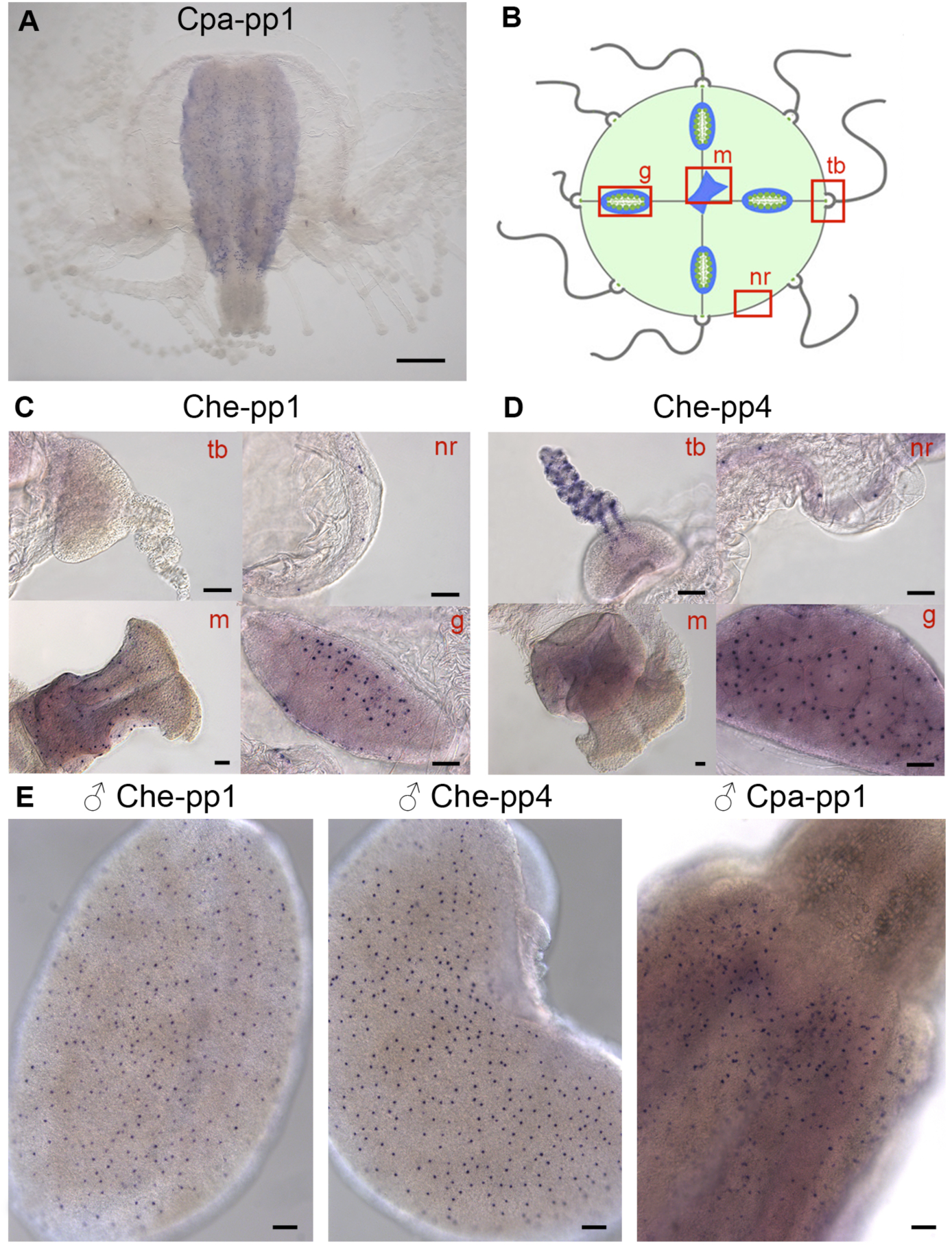
Figure 5. Distribution of MIH-expressing cells detected by *in situ* hybridisation. A) Cpa-pp1 in scattered ectodermal cells in the gonad of a *Cladonema* female jellyfish. B) Schematic representation of a *Clytia* jellyfish indicating the position of the tentacle bulbs (tb), nerve ring (nr), gonads (g) and manubrium (m). C-D) *In situ* hybridisation detection of Che-pp1 and Che-pp4 in different structures of young *Clytia* jellyfish as indicated in red. Both these precursors are expressed in the gonad ectoderm and nerve ring, but in the manubrium mainly Che-pp1 is detected, and in the tentacle Che-pp4. E) *In situ* hybridisation detection in male gonads from *Clytia* and *Cladonema* showing Che-pp1, Che-pp4 and Cpa-pp1 expression in scattered ectoderm cells. Scale bars: 500 *μ*m in A; 50 *μ*m in C-E.

In *Clytia* gonad ectoderm, the anti-PRPamide and PRAamide antibodies decorated a single cell population, whereas in *Cladonema* the two peptides were detected in distinct cell populations (Fig. S5B, C), presumably being generated from the Cpa-PP1 and Cpa-PP2 precursors, respectively (Fig. S1). Immunofluorescence analysis of *Cladonema* gonads found a reduction of the anti-PRPamide signal within 20 minutes after darkness, whereas the anti-PRAamide signal was relatively unaffected (Fig. 4D). *Clytia* gonads showed a moderate reduction of staining with both antibodies 45 and 120 minutes after light stimulus (Fig. 4E). More detailed examination indicated that in each stained cell the numbers of antibody-positive dots, presumably representing peptide-filled vesicles, decreased following light exposure in both the cell body and in the microtubule-rich projections (Fig. 4F). It would be interesting to determine, for instance by live imaging techniques, whether MIH-containing vesicles are secreted from particular sites or exhibit any trafficking following the light/dark cues, or whether indeed vesicle release occurs throughout the cell, as would be typical of a neurosecretory cell type (Hartenstein, 2006).

The similar distribution of MIH-producing cells in female and male gonads (Fig. 4A, 5E) suggests that these neuropeptides may play a general role in regulating gamete release, and not only in the initiation of oocyte maturation in female medusae. We found using male jellyfish of both *Clytia* and *Cladonema* that synthetic MIH peptides at 10^−7^ M provoked release of active sperm from the gonads (Table 1). This finding confirms that the oocyte maturation stimulating effect of MIH is part of a wider role in reproductive regulation. It also raises the intriguing possibility that MIH neuropeptides released into the seawater from males and females gathered together at the ocean surface during spawning may facilitate precise synchronisation of gamete release during the periods of dawn and dusk.

**Table 1.**
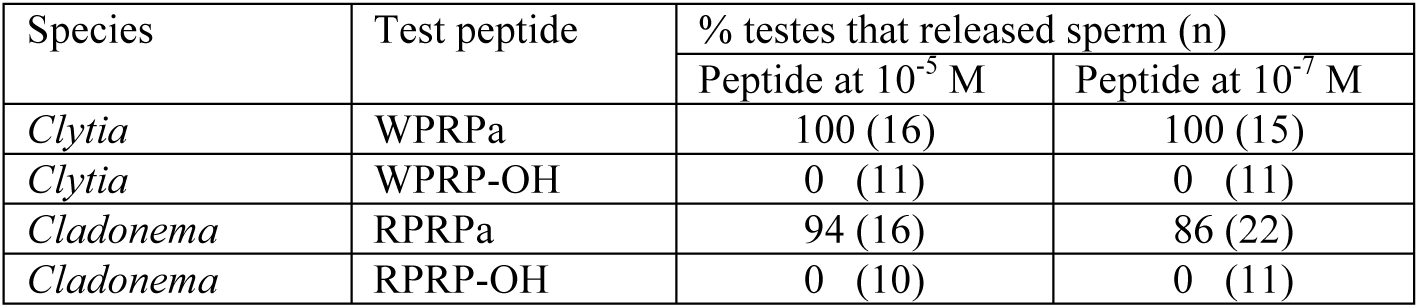
MIH peptides induce male spawning.

### Selective action of MIH peptides between hydrozoan jellyfish species

Our experiments revealed some selectivity in the MIH activity of different peptides between *Clytia* and *Cladonema*. The most potent MIH peptides for *Clytia* oocytes were the main Che-pp1/Che-pp4-derived tetrapeptides WPRPamide, WPRAamide and RPRYamide, clearly active even at 10^−8^ M (Fig. S3). The best candidate for *Cladonema* MIH is RPRPamide (from Cpa-pp1), while RPRAamide (from Cpa-pp2) was slightly less active (Fig. S3). Correspondingly, the RPRP sequence is not found in precursors expressed in the *Clytia* gonad, while WPRP/Aamides are not predicted from any *Cladonema* precursors (see Fig. 2A; Fig. S1). Further testing on oocytes from eight other hydrozoan jellyfish species revealed responsiveness with different sensitivities to W/RPRP/A/G/Yamide type tetrapeptides in *Obelia, Aequorea, Bouillonactinia* and *Sarsia*, but not *Eutonina, Nemopsis, Rathkea* or *Cytaeis* (Fig. 6). The responsive and non-responsive species included members of two main hydrozoan groups, leptomedusae and anthomedusae. These comparisons suggest that W/RPRXamide type peptides functioned as MIHs in ancestral hydrozoan jellyfish. We can speculate that variation in the peptide sequences active between related species might reduce stimulation of spawning between species in mixed wild populations.

**Figure 6.**
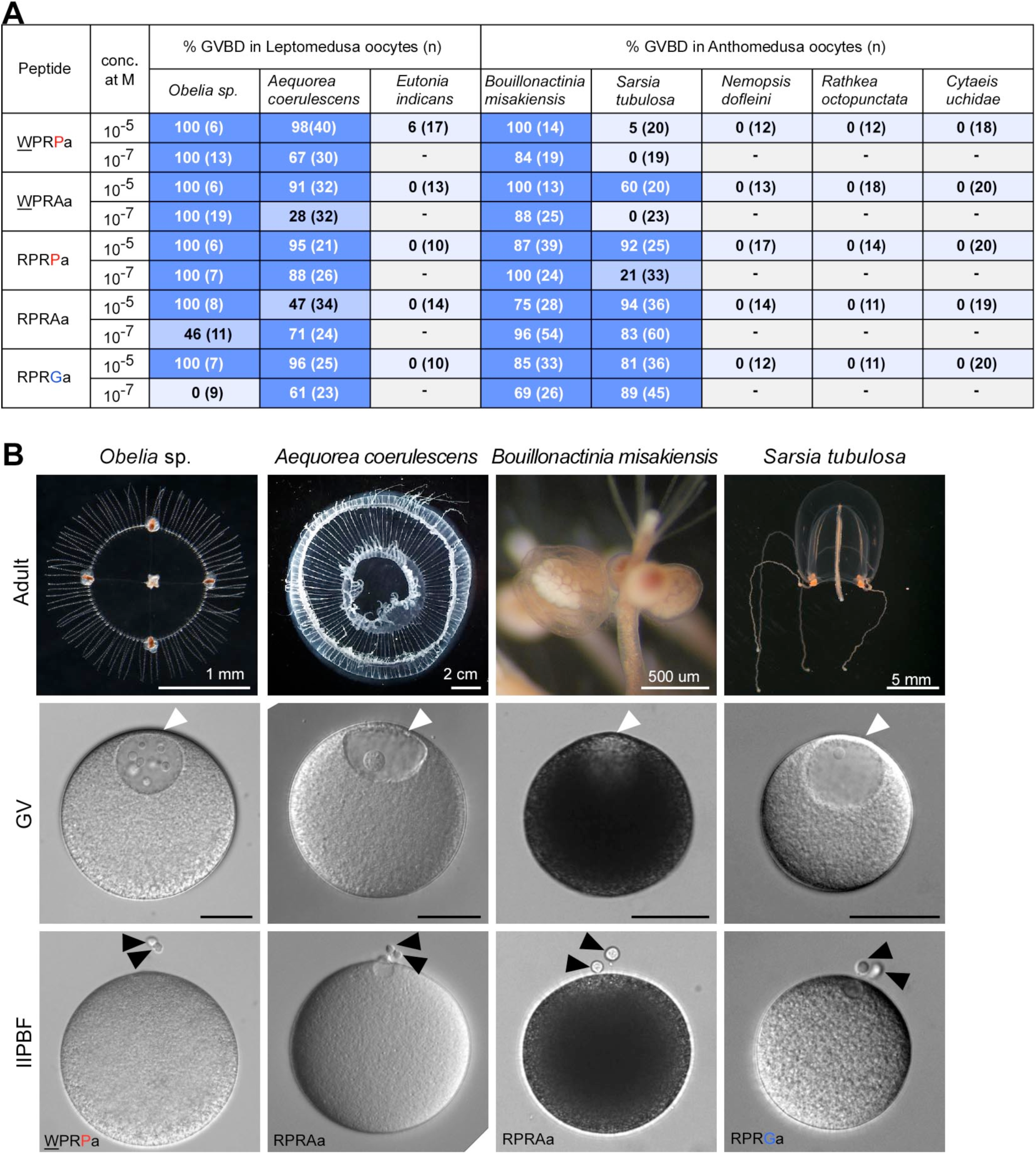
Synthetic peptides show MIH activity in a subset of hydrozoan jellyfish species. A) Synthetic W/RPRP/A/Gamides were tested for their ability at concentrations of 10^−5^ M and 10^−7^ M to induce GVBD of oocytes of the 8 species indicated. Highest success of GVBD is emphasised by the darker blue colours. B) Examples of four of the species tested, showing the adult females (top row), isolated oocytes (middle row) and mature eggs with two polar bodies (bottom row) generated by incubation in the peptides indicated. Scale bars for oocytes: 50μm. White and black arrowheads indicate GVs and polar bodies, respectively.

### Discussion

We have demonstrated that short amidated peptides with the prototype sequence W/RPRPamide are responsible for inducing oocyte maturation, resulting ultimately release from the gonad of active mature gametes in the hydrozoan jellyfish *Clytia* and *Cladonema* (Fig. 7A). These peptides act as *bona fide* MIHs, i.e. they interact directly with the surface of resting ovarian oocytes to initiate maturation. Related W/RPRXamide peptides act as MIHs also in other hydrozoan jellyfish species. Concerning the later events of the spawning process, we hypothesise that egg release through the gonad ectoderm in female medusae is not triggered directly by MIH but via a secondary signal emitted by the oocyte towards the end of the meiotic maturation process. More specifically it could depend on Mos-MAP kinase activation in the oocyte, since maturation in the absence of spawning can occur when Mos translation is inhibited in ovarian oocytes (Amiel et al., 2009). In male gonads MIH peptides presumably act on inactive, post-meiotic spermatozoids to initiate the spawning response but the mechanisms involved are not yet known.

**Figure 7.**
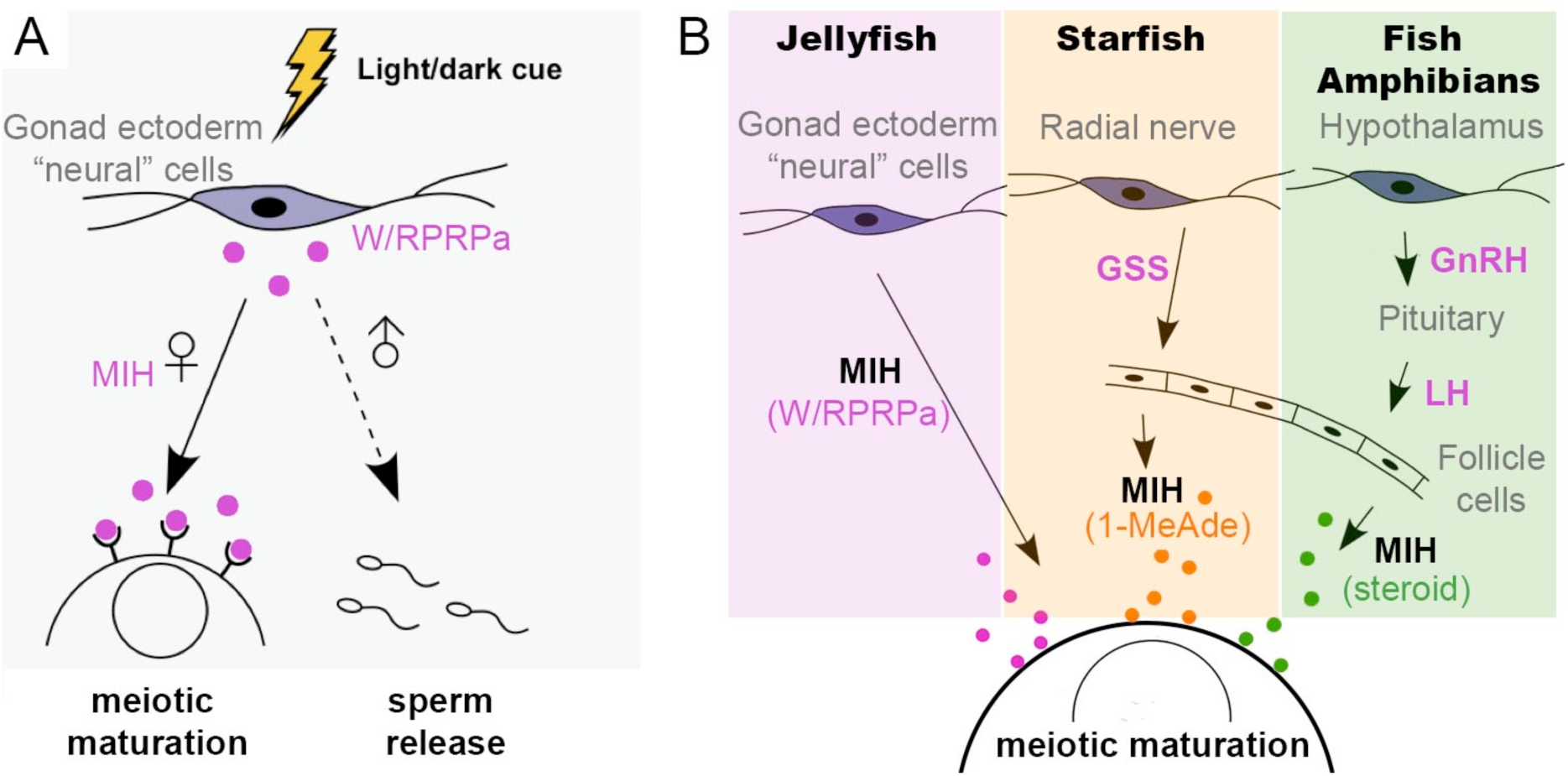
MIH action in jellyfish compared with other animals. A) Summary of the findings of this study: jellyfish MIH, consisting of PRPamide family peptides in *Clytia* and *Cladonema*, is secreted by neural-type cells in the gonad directly in response to light cues and causes oocyte maturation as well as spawning in males and females. B) Comparison of the regulation of oocyte maturation by peptide hormones (pink) in jellyfish, starfish and fish/amphibians.

Some GLWamide family peptides are also able to provoke oocyte maturation and spawning, albeit at higher concentrations than the PRPamides, but do not induce maturation of isolated oocytes (this study and Takeda et al., 2013), suggesting that the role of these peptides in regulating spawning is indirect. We can imagine that inhibitory or sensitising factors, possibly including GLWamides, could act either in the gonad MIH-secreting cells or in other ectodermal cells to modulate the light response, and account for species-specific dawn or dusk spawning. It remains to be seen whether regulation of spawning by MIH neuropeptides related to those in *Clytia* and *Cladonema* extends beyond hydrozoan jellyfish to other cnidarians. If so, further layers of regulation could allow the integration of seasonal cues and lunar cycles to account for well known mass annual spawning events seen in tropical reef corals (Harrison et al., 1984).

The identification of MIH in *Clytia* and *Cladonema* is a significant step forward in the oocyte maturation field because the molecular nature of the hormones that in this way trigger oocyte maturation is known in surprisingly few animal species, notably 1-methyladenine in starfish and steroid hormones in teleost fish and amphibians (Kanatani et al., 1969; Nagahama et al., 2008; Yamashita et al., 2000; Haccard et al., 2012). The very different molecular natures of these known MIH examples from across the (bilaterian+cnidarian) clade could be explained by an evolutionary scenario in which secretion of neuropeptide MIHs from cells positioned near to the oocyte was the ancestral condition, with intermediate regulatory tissues, such as endocrine organs and ovarian follicle cells, evolving in the deuterostome lineage to separate neuropeptide-based regulation from the final response of the oocyte (Fig. 7B). Such interpolation of additional layers of regulation is a common feature of endocrine system evolution (Hartenstein, 2006). During the evolution of reproductive regulation, various neuropeptides including vertebrate GnRHs (gonadotropin-releasing hormones; (Roch et al., 2011), as well as modulatory RFamide peptides such as Kisspeptins and GnIH (gonadotropin-inhibitory hormone; Parhar et al., 2012), regulate various aspects of reproduction including gamete release in both males and females. Chordate GnRHs are PGamide decapeptides, which stimulate the release of peptidic gonadotropic hormones (GTHs) such as vertebrate luteinizing hormone from the pituitary. Similarly, starfish gonad-stimulating substance (GSS/Relaxin; Mita et al., 2009) is a GTH produced at a distant “neuroendocrine” site, the radial nerve. In both cases, these peptidic GTHs in turn cause oocyte maturation by inducing MIH release from the surrounding follicle cells, or in the case of mammals GAP junction-mediated exchange of cyclic nucleotides between these cells (Shuhaibar et al., 2015). Regulation of reproduction by GnRHs probably predated the divergence of deuterostomes and protostomes (Roch et al., 2011; Tsai, 2006), the best evidence coming from mollusc species in which peptides structurally related to GnRH, synthesised at various neuroendocrine sites, regulate various reproductive processes (Osada and Treen, 2013).

Cnidarians use neuropeptides to regulate multiple processes including muscle contraction, neural differentiation and metamorphosis from larva to polyp (Anctil, 2009; Fujisawa, 2008; Takahashi and Hatta, 2011). Transcript sequences predicted to produce many copies of short neuropeptides have also been found in ctenophore and placozoan genomes (Moroz et al., 2014; Nikitin, 2015), and neuropeptides are thought to have been the predominant neurotransmitters in the ancient common ancestor of these groups (Grimmelikhuijzen and Hauser, 2012). Although independent evolution of neuropeptide regulation or reproduction between animal clades cannot be ruled out, the identification of the MIH neuropeptides in *Clytia and Cladonema* along with other evidence from cnidarians (Takeda et al., 2013; Tremblay et al., 2004) as well as bilaterians (see above), suggests that neuropeptide signalling played a central role in coordinating sexual reproduction in the bilaterian-cnidarian ancestor, and may have been involved in coordinating spawning events in the marine environment. In *Clytia* medusae we found cells producing PRPamide family peptides not only in the gonad but also in the manubrium, tentacles and bell margin (Fig. 5C,D), so presumably these have wider functions than orchestrating gamete release. It will be of great interest to investigate the activities of related peptides across a wide range of species in order to track the evolutionary history of the neuroendocrine regulation of reproduction.

## Materials and Methods

### Animal cultures

Laboratory strains of *Clytia hemisphaerica* (“Z colonies”), *Cladonema pacificum* (6W, NON5, UN2), and *Cytaeis uchidae* (17) were maintained throughout the year (Deguchi et al., 2005; Houliston et al., 2010; Takeda et al., 2006). Wild specimens of *Cladonema* as well as *Eutonina indicans, Nemopsis dofleini, Obelia* sp., *Rathkea octopunctata*, and *Sarsia tublosa* were collected from Sendai Bay, Miyagi Prefecture, and *Aequorea coerulescens* and *Bouillonactinia misakiensis* from Mutsu Bay, Aomori Prefecture. The brand of artificial seawater (ASW) used for culture and for functional assays in Japan was SEA LIFE (Marine Tech, Tokyo), and for *Clytia hemisphaerica* culture, transcriptomics and microscopy in France was Red Sea Salt.

### Oocyte isolation and MIH assays

Fully-grown oocytes were obtained from ovaries of intact jellyfish or pre-isolated ovaries placed under constant illumination for 20-24 h following the previous spawning. Ovarian oocytes were aspirated using a mouth pipette or detached using fine tungsten needles. During oocyte isolation, jellyfish were in some cases anaesthetised in excess Mg^2+^ ASW (a 1:1 mix of 0.53 M MgCl_2_ and ASW). Pre-isolated ovaries of *Clytia, Aequorea*, and *Eutonina* were bathed in ASW containing 1 mM sodium citrate to facilitate the detachment of oocytes from ovarian tissues.

Active MIH was recovered from cultured ovaries of *Clytia* and *Cladonema* by a similar approach to that used previously (Freeman, 1987). A chamber formed between a plastic dish and a coverslip separated by two pieces of 400 or 500 *μ*m-thick double-stick tape was filled with silicon oil (10 cSt; TSF451-10, Momentive Performance Materials), and a drop of ASW (0.5-1 *μ*L) containing several ovaries separated from 2 *Clytia* jellyfish or several ovarian epithelium fragments stripped from 3-5 *Cladonema* jellyfish were inserted into the oil space (Fig. 1A,B). The oil chambers were subjected to light-dark changes (light after dark in *Clytia* and dark after light in *Cladonema*) and the ASW with MIH activity was collected 60 min later. Prior to MIH assays, isolated oocytes were cultured in seawater for at least 30 min and any oocytes showing damage or GVBD discarded. MIH assays were performed at 14-16°C for *Eutonina, Nemopsis, Obelia, Sarsia* and *Rathkea* or 18-21°C for *Clytia, Cladonema, Cytaeis*, and *Bouillonactinia*.

### Identification of peptide precursors

Potential amidated peptide precursor sequences were recovered from a *Clytia* reference transcriptome derived from mixed larva, polyp and jellyfish samples. ORFs and protein sequences were predicted using an R script (Lapébie et al., 2014). Potential secreted proteins were identified by the presence of signal peptide, using SignalP 4.0 (Petersen et al., 2011). Then sequences rich in the amidated pro-peptide cleavage motifs GR/K and lacking domains recognised by Interproscan-5.14-53.0 were selected. Finally, sequences containing repetitive motifs of less than 20 amino-acids were identified using TRUST (Szklarczyk and Heringa, 2004). Among this final set of putative peptide precursors, some known secreted proteins with repetitive structures were identified by BLAST and removed.

To prepare a *Cladonema* transcriptome, more than 10 *μ*g of total RNA was isolated from the manubrium of female jellyfish (6W strain) using NucleoSpin RNA purification kit (MACHEREY-NAGEL, KG). RNA-seq library preparation and sequencing Ilumina HiSeq 2000) were carried out by BGI (Hong-Kong, China). Using an assembled dataset containing 74,711 contigs and 35,957 unigenes, local BLAST searches were performed to find peptide precursors using published cnidarian neuropeptide sequences or the *Clytia* pp1 and pp4 sequences as bait.

The ORFs of putative candidate *Clytia* and *Cladonema* peptide precursors were cloned by PCR into pGEM-T easy vector, or retrieved from our *Clytia* EST collection cDNA library prior to probe synthesis. Sequences and accession numbers are given in Table S1.

For *Clytia* gonad tissue transcriptome comparisons, Illlumina Hi-seq 50nt reads were generated from mRNA isolated using RNAqueous micro kit (Ambion Life technologies, CA) from ectoderm, endoderm and oocytes manually dissected from about 150 *Clytia* female gonads. Q-PCR was performed to check for contamination between samples using endogenous GFP genes expressed in oocyte, ectoderm and bell tissue (Fourrage et al., 2014), and to quantify expression of selected peptide precursors (primer list in Fig. S2B). The reads were mapped against a *Clytia* reference transcriptome using Bowtie2 (Langmead and Salzberg, 2012). The counts for each contig were normalised per total of reads of each sample and per sequence length and visualised using the heatmap.2 function in the “gplots” R package.

### Peptides and antibodies

WPRP-NH_2_, WPRA-NH_2_, RPRP-NH_2_, RPRA-NH_2_, RPRG-NH_2_, RPRY-NH_2_, PGLW-NH_2_, DAWPRP-NH_2_, AWPRP-NH_2_, FNIRPRP-NH_2_, NIRPRP-NH_2_, IRPRP-NH_2_, PRP-NH_2_,WPRP-OH and RPRP-OH were synthesised by GenScript or Life Technologies. These peptides were dissolved in deionised water at 10^−2^ M or 2x10^−3^ M, stored at −20°C, and diluted in ASW at 10^−5^–10^−10^ M prior to use. TAMRA-WPRPamide (TAMRA-LEKRNWPRP-NH_2_); was synthesised by Sigma and a 5x10^−4^ M solution in H_2_O was injected at 2–17% of the oocyte volume, to give an estimated final oocyte concentration of 1 to 9x10^−5^ M (Deguchi et al., 2005).

Polyclonal antibodies against XPRPamide and XPRAamide were raised in rabbits using keyhole limpet hemocyanin (KLH)-conjugated CPRA-NH_2_ and CPRP-NH_2_ as antigens, and antigen-specific affinity purified (Sigma-Ardrich Japan). For MIH inhibition experiments, antibodies or control normal rabbit IgG (MBL) were concentrated using a 30000 MW cut-off membrane (Millipore), giving a final protein concentration of 10^−6^ M, and the buffers were replaced with seawater through repeated centrifugations.

### Immunofluorescence and in situ hybridisation

For single or double anti-PRPamide /anti-PRAamide staining, specimens were pre-anesthetised using excess Mg^2+^ ASW and fixed overnight at 4°C in 10% formalin-containing ASW and rinsed 3x 10 min in Phosphate Buffered Saline (PBS) containing 0.25% Triton X-100. They were incubated in anti-PRPamide or anti-PRAamide antibody diluted in PBSTriton overnight at 4°C. After washes in PBS-Triton, the specimens were incubated with secondary antibody (Alexa Fluor 488 or 568 goat anti-rabbit IgG; Invitrogen, Carlsbad, CA) for 2 h at room temperature and nuclei stained using 50 *μ*M Hoechst 33258 or 33342 (Invitrogen) for 5-20 min. Zenon antibody labeling kits (Molecular Probes, Eugene, OR) were used for double peptide staining. In control experiments, PBS-Triton alone or normal rabbit IgG (3 mg/ml; Zymed, San Francisco, CA) in PBS-Triton (1/1000 dilution) replaced the anti-PRPamide or anti-PRAamide antibodies. Images were acquired using a laser scanning confocal system (C1, Nikon).

For co-staining of neuropeptides and microtubules (Fig. 4C,D), dissected *Clytia* gonads were fixed overnight at 18°C in HEM buffer (0.1 M HEPES pH 6.9, 50 mM EGTA, 10 mM MgSO_4_) containing 3.7% formaldehyde, then washed five times in PBS containing 0.1% Tween20 (PBS-T). Treatment on ice with 50% methanol/PBS-T then 100% methanol plus storage in methanol at −20°C improved visualisation of microtubules in the MIH-producing cells. Samples were rehydrated, washed in PBS-0.02% Triton X-100, blocked in PBS with 3% BSA overnight at 4°C, then incubated in anti-PRPamide antibody and anti-alpha tubulin (YL1/2) in PBS/BSA at room temperature for 2 h. After washes, the specimens were incubated with secondary antibodies (Rhodamine goat anti-rabbit and Cy5 donkey anti-rat-IgG; Jackson ImmunoResearch, West Grove, PA) overnight in PBS at 4°C and nuclei stained using Hoechst dye 33258 for 20 min.

For *in situ* hybridisation, isolated gonads or whole jellyfish were processed as previously (Fourrage et al., 2014) except that 4 M Urea was used instead of 50% formamide in the hybridisation buffer (Sinigaglia et al., 2017). For double fluorescent *in situ* hybridisation, female *Clytia* gonads were fixed overnight at 18°C in HEM buffer containing 3.7% formaldehyde, washed five times in PBS containing 0.1% Tween20 (PBS-T), then dehydrated on ice using 50% methanol/PBS-T then 100% methanol. *In situ* hybridisation (Lapébie et al., 2014) (Sinigaglia et al., 2017) was performed using a DIG-labeled probe for Che-pp1 and a fluorescein-labeled probe for Che-pp4. A 3 hour incubation with a peroxidase labeled anti-DIG antibody was followed by washes in MABT (100 mM maleic acid pH 7.5, 150 mM NaCl, 0.1% Triton X-100). For Che-pp1 the fluorescence signal was developed using the TSA (Tyramide Signal Amplification) kit (TSA Plus Fluorescence Amplification kit, PerkinElmer, Waltham, MA) and Cy3 fluorophore (diluted 1/400 in TSA buffer: PBS/H_2_O_2_ 0.0015%) at room temperature for 30 min. After 3 washes in PBS-T fluorescence was quenched with 0.01 M HCl for 10 min at room temperature and washed again several times in PBS-T. Overnight incubation with a peroxidase-labelled anti-fluorescein antibody was followed by washes in MABT. The anti Che-pp4 fluorescence signal was developed using TSA kit with Cy5 fluorophore. Nuclei were stained using Hoechst dye 33258. Images were acquired using a Leica SP5 confocal microscope and maximum intensity projections of z-stacks prepared using ImageJ software.

## Acknowledgements

We thank P. Dru S. Chevalier and L. Leclère for generating and assembling *Clytia* reference transcriptome, A. Ruggiero and C. Sinigaglia for sharing *in situ* hybridisation protocols, S. Yaguchi for useful advice on immunofluorescence and J. Uveira for technical assistance. We also thank our group members, “Neptune” network colleagues, Clare Hudson and Hitoyoshi Yasuo for useful discussions. Work was supported by JSPS KAKENHI Grant Numbers 26440177 & 26840073, the French ANR (“OOCAMP”) grant, the Marie Curie ITN “Neptune” and the Tokyo Institute of Technology GCOE program from JSPS (NT’s visit to Villefranche). Microscopy equipment at the Villefranche-sur-mer imaging platform was cofinanced by the PACA region, CNRS and UPMC.

## References

Amiel, A., Chang, P., Momose, T. and Houliston, E. (2010). *Clytia hemisphaerica:* a cnidarian model for studying oogenesis. In Oogenesis: the universal process. Chichester: John Wiley & Sons. 81–102.

Amiel, A., Leclère, L., Robert, L., Chevalier, S. and Houliston, E. (2009). Conserved functions for Mos in eumetazoan oocyte maturation revealed by studies in a cnidarian. Curr. Biol. 19, 305–311.

Anctil, M. (2000). Evidence for Gonadotropin-Releasing Hormone-like peptides in a cnidarian nervous system. Gen. Comp. Endocrinol. 119, 317–328.

Anctil, M. (2009). Chemical transmission in the sea anemone *Nematostella vectensis:* A genomic perspective. Comparative Biochemistry and Physiology - Part D: Genomics and Proteomics 4, 268–289.

Bosch, T. C. G., Klimovich, A., Domazet-Lošo, T., Gründer, S., Holstein, T. W., Jékely, G., Miller, D. J., Murillo-Rincon, A. P., Rentzsch, F., Richards, G. S., et al. (2017). Back to the Basics: Cnidarians Start to Fire. Trends in Neurosciences 40, 92–105.

David, C. N. (1973). A quantitative method for maceration of hydra tissue. Wilhelm Roux Arch Entwickl Mech Org 171, 259–268.

Deguchi, R., Kondoh, E. and Itoh, J. (2005). Spatiotemporal characteristics and mechanisms of intracellular Ca^2^+ increases at fertilization in eggs of jellyfish (Phylum Cnidaria, Class Hydrozoa). Dev. Biol. 279, 291–307.

Dupre, C. and Yuste, R. (2017). Non-overlapping Neural Networks in *Hydra vulgaris*. Curr. Biol. 27, 1085–1097.

Fourrage, C., Swann, K., Gonzalez Garcia, J. R., Campbell, A. K. and Houliston, E. (2014). An endogenous green fluorescent protein-photoprotein pair in *Clytia hemisphaerica* eggs shows co-targeting to mitochondria and efficient bioluminescence energy transfer. Open Biol 4, 130206.

Freeman, G. (1987). The role of oocyte maturation in the ontogeny of the fertilization site in the hydrozoan *Hydractinia echinata*. Roux’s Arch Dev. Biol. 196, 83–92.

Fujisawa, T. (2008). Hydra peptide project 1993-2007. Dev. Growth Differ. 50 Suppl 1, S257–68.

Grimmelikhuijzen, C. J. P. and Hauser, F. (2012). Mini-review: the evolution of neuropeptide signaling. Regul. Pept. 177 Suppl, S6–9.

Grimmelikhuijzen, C. J. P., Leviev, I. and Carstensen, K. (1996). Peptides in the nervous systems of cnidarians: Structure, function and biosynthesis. In International Review of Cytology, pp. 37–89. Elsevier.

Gründer, S. and Assmann, M. (2015). Peptide-gated ion channels and the simple nervous system of Hydra. Journal of Experimental Biology 218, 551–561.

Haccard, O., Dupré, A., Liere, P., Pianos, A., Eychenne, B., Jessus, C. and Ozon, R. (2012). Naturally occurring steroids in *Xenopus* oocyte during meiotic maturation. Unexpected presence and role of steroid sulfates. Mol. Cell. Endocrinol. 362, 110–119.

Hartenstein, V. (2006). The neuroendocrine system of invertebrates: a developmental and evolutionary perspective. Journal of Endocrinology 190, 555–570.

Harrison, P. L., Babcock R. C., Bull, G. D., Oliver, J. K., Wallace C. C. and Willis B. L. (1984). Mass spawning in tropical reef corals. Science 223, 1186–1189.

Houliston, E., Momose, T. and Manuel, M. (2010). *Clytia hemisphaerica:* a jellyfish cousin joins the laboratory. Trends Genet. 26, 159–167.

Ikegami, S., Honji, N. and Yoshida, M. (1987). Light-controlled production of spawning-inducing substance in jellyfish ovary. Nature 272, 611–612.

Kanatani, H., Shirai, H., Nakanishi, K. and Kurokawa, T. (1969). Isolation and indentification on meiosis inducing substance in starfish *Asterias amurensis*. Nature 221, 273–274.

Koizumi, O., Hamada, S., Minobe, S., Hamaguchi-Hamada, K., Kurumata-Shigeto, M., Nakamura, M. and Namikawa, H. (2015). The nerve ring in cnidarians: its presence and structure in hydrozoan medusae. Zoology 118, 79–88.

Koizumi, O. (2016). Origin and Evolution of the Nervous System Considered from the Diffuse Nervous System of Cnidarians. In The Cnidaria, Past, Present and Future, pp. 73–91. Cham: Springer International.

Langmead, B. and Salzberg, S. L. (2012). Fast gapped-read alignment with Bowtie 2. Nat. Methods 9, 357–359.

Lapébie, P., Ruggiero, A., Barreau, C., Chevalier, S., Chang, P., Dru, P., Houliston, E. and Momose, T. (2014). Differential responses to Wnt and PCP disruption predict expression and developmental function of conserved and novel genes in a cnidarian. PLoS Genet 10, e1004590.

Mita, M., Yoshikuni, M., Ohno, K., Shibata, Y., Paul-Prasanth, B., Pitchayawasin, S., Isobe, M. and Nagahama, Y. (2009). A relaxin-like peptide purified from radial nerves induces oocyte maturation and ovulation in the starfish *Asterina pectinifera*. Proc. Natl. Acad. Sci. U.S.A. 106, 9507–9512.

Moroz, L. L., Kocot, K. M., Citarella, M. R., Dosung, S., Norekian, T. P., Povolotskaya, I. S., Grigorenko, A. P., Dailey, C., Berezikov, E., Buckley, K. M., et al. (2014). The ctenophore genome and the evolutionary origins of neural systems. Nature 510, 109–114.

Nagahama, Y. and Yamashita, M. (2008). Regulation of oocyte maturation in fish. Dev. Growth Differ. 50 Suppl 1, S195–219.

Nikitin, M. (2015). Bioinformatic prediction of trichoplax adhaerens regulatory peptides. Gen. Comp. Endocrinol. 212, 145–155.

Osada, M. and Treen, N. (2013). Molluscan GnRH associated with reproduction. Gen. Comp. Endocrinol. 181, 254–258.

Parhar, I., Ogawa, S. and Kitahashi, T. (2012). RFamide peptides as mediators in environmental control of GnRH neurons. Progress in Neurobiology 98, 176–196.

Petersen, T. N., Brunak, S., Heijne von, G. and Nielsen, H. (2011). SignalP 4.0: discriminating signal peptides from transmembrane regions. Nat. Methods 8, 785–786.

Quiroga Artigas, G., Lapébie, P., Leclère, L., Takeda, N., Deguchi, R., Jékely, G., Momose, T. and Houliston, E. (2017). CRISPR/Cas9 mutation of a gonad-expressed opsin prevents jellyfish light-induced spawning. http://biorxiv.org/lookup/doi/10.1101/140210

Roch, G. J., Busby, E. R. and Sherwood, N. M. (2011). Evolution of GnRH: Diving deeper. Gen. Comp. Endocrinol. 171, 1–16.

Shuhaibar, L. C., Egbert, J. R., Norris, R. P., Lampe, P. D., Nikolaev, V. O., Thunemann, M., Wen, L., Feil, R. and Jaffe, L. A. (2015). Intercellular signaling via cyclic GMP diffusion through gap junctions restarts meiosis in mouse ovarian follicles. Proc. Natl. Acad. Sci. U.S.A. 112, 5527–5532.

Sinigaglia, C., Thiel, D., Hejnol, A., Houliston, E. and Leclère, L. (2017). A safer, urea-based in situ hybridization method improves detection of gene expression in diverse animal species. http://biorxiv.org/lookup/doi/10.1101/133470

Szklarczyk, R. and Heringa, J. (2004). Tracking repeats using significance and transitivity. Bioinformatics 20 Suppl 1, i311–7.

Tachibana, K., Tanaka, D., Isobe, T. and Kishimoto, T. (2000). c-Mos forces the mitotic cell cycle to undergo meiosis II to produce haploid gametes. Proc. Natl. Acad. Sci. U.S.A. 97, 14301–14306.

Takahashi, T. and Hatta, M. (2011). The importance of GLWamide neuropeptides in cnidarian development and physiology. J Amino Acids 2011, 424501.

Takahashi, T., Hayakawa, E., Koizumi, O. and Fujisawa, T. (2008). Neuropeptides and their functions in *Hydra*. Acta. Biol. Hung. 59 Suppl, 227–235.

Takeda, N., Kyozuka, K. and Deguchi, R. (2006). Increase in intracellular cAMP is a prerequisite signal for initiation of physiological oocyte meiotic maturation in the hydrozoan *Cytaeis uchidae*. Dev. Biol. 298, 248–258.

Takeda, N., Nakajima, Y., Koizumi, O., Fujisawa, T., Takahashi, T., Matsumoto, M. and Deguchi, R. (2013). Neuropeptides trigger oocyte maturation and subsequent spawning in the hydrozoan jellyfish *Cytaeis uchidae*. Mol. Reprod. Dev. 80, 223–232.

Tremblay, M.-E., Henry, J. and Anctil, M. (2004). Spawning and gamete follicle rupture in the cnidarian *Renilla koellikeri:* effects of putative neurohormones. Gen. Comp. Endocrinol. 137, 9–18.

Tsai, P.-S. (2006). Gonadotropin-releasing hormone in invertebrates: Structure, function, and evolution. Gen. Comp. Endocrinol. 148, 48–53.

Von Stetina, J. R. and Orr-Weaver, T. L. (2011). Developmental control of oocyte maturation and egg activation in metazoan models. Cold Spring Harb Perspect Biol 3, a005553.

Yamashita, M., Mita, K., Yoshida, N. and Kondo, T. (2000). Molecular mechanisms of the initiation of oocyte maturation: general and species-specific aspects. Prog Cell Cycle Res 4, 115–129.

